# A COVID-19 antibody curbs SARS-CoV-2 nucleocapsid protein-induced complement hyper-activation

**DOI:** 10.1101/2020.09.10.292318

**Authors:** Sisi Kang, Mei Yang, Suhua He, Yueming Wang, Xiaoxue Chen, Yao-Qing Chen, Zhongsi Hong, Jing Liu, Guanmin Jiang, Qiuyue Chen, Ziliang Zhou, Zhechong Zhou, Zhaoxia Huang, Xi Huang, Huanhuan He, Weihong Zheng, Hua-Xin Liao, Fei Xiao, Hong Shan, Shoudeng Chen

## Abstract

Although human antibodies elicited by severe acute respiratory distress syndrome coronavirus-2 (SARS-CoV-2) nucleocapsid (N) protein are profoundly boosted upon infection, little is known about the function of N-directed antibodies. Herein, we isolated and profiled a panel of 32 N protein-specific monoclonal antibodies (mAb) from a quick recovery coronavirus disease-19 (COVID-19) convalescent, who had dominant antibody responses to SARS-CoV-2 N protein rather than to Spike protein. The complex structure of N protein RNA binding domain with the highest binding affinity mAb nCoV396 reveals the epitopes and antigen’s allosteric changes. Functionally, a virus-free complement hyper-activation analysis demonstrates that nCoV396 specifically compromises N protein-induced complement hyper-activation, a risk factor for morbidity and mortality in COVID-19, thus paving the way for functional anti-N mAbs identification.

**One Sentence Summary:** B cell profiling, structural determination, and protease activity assays identify a functional antibody to N protein.

## Main Text

The fatality rate of the critical condition <u>Co</u>rona<u>vi</u>rus <u>D</u>isease 2019 (COVID-19) patients is exceptionally high, at 40% - 49%(*1, 2*). Acute respiratory failure and generalized coagulopathy are significant aspects associated with morbidity and mortality(*3–5*). A subset of severe COVID-19 patients has distinct clinical features compared to classic <u>a</u>cute <u>r</u>espiratory <u>d</u>istress <u>s</u>yndrome (ARDS), with delayed onset of respiratory distress(*6*) and relatively well-preserved lung mechanics despite the severity of hypoxemia(*7*). It is reported that complement-mediated thrombotic microvascular injury in the lung may contribute to atypical ARDS features of COVID-19, accompanied by extensive deposition of the <u>a</u>lternative <u>p</u>athway (AP) and <u>l</u>ectin <u>p</u>athway (LP) complement components(*8*). Indeed, complement activation is found in multiple organs of severe COVID-19 patients in several other studies(*9, 10*), as well as in patients with <u>s</u>evere <u>a</u>cute <u>r</u>espiratory distress <u>s</u>yndrome (SARS)(*11, 12*). A recent retrospective observational study of 11,116 patients revealed that complement disorder associated with morbidity and mortality of COVID-19(*13*).

Although systemic activation of complement plays a pivotal role in protective immunity against pathogens, hyper-activation of complement may lead to collateral tissue injury. <u>S</u>evere <u>a</u>cute <u>r</u>espiratory distress <u>s</u>yndrome-associated <u>co</u>rona<u>v</u>irus-<u>2</u> (SARS-CoV-2) nucleocapsid (N) protein is a highly immunopathogenic and multifunctional viral protein*(14–19)*, which elicited high titers of binding antibodies in humoral immune responses(*20–22*). A recent preprint study found that SARS-CoV-2 N protein bound to LP complement components MASP-2 (<u>M</u>annan binding lectin-<u>a</u>ssociated serine <u>p</u>rotease-<u>2</u>), and resulted in complement hyper-activation and aggravated inflammatory lung injury(*15*). Several studies have reported in isolations of human <u>m</u>onoclonal <u>a</u>nti<u>b</u>odie<u>s</u> (mAbs) targeting SARS-CoV-2 Spike (S) protein, shedding the light of developing therapeutic interventions of COVID-19(*20, 23–27*). However, little is known about the potential therapeutic applications of N protein-targeting mAbs in the convalescent B cell repertoire. Herein, we report a human mAb derived from COVID-19 convalescent, with specific targeting to SARS-CoV-2 N protein and functionally compromising complement hyper-activation *ex vivo*.

### Isolation of N protein-directed mAbs

To profile antibody response to SARS-CoV-2 N protein in early recovered patients, we collected six convalescent blood samples at seven to 25 days after the onset of the disease symptoms. All patients are recovered from COVID-19 during the outbreak in Zhuhai, Guangdong Province, China, with age ranging from 23 to 66 years old (**Table S1**). The SARS-CoV-2 nasal swabs reverse transcription-polymerase chain reaction (RT-PCR) tests were confirmed being negative at the points of blood collection for all of these six COVID-19 patients. Plasma samples and peripheral blood mononuclear cells (PBMC) were isolated for serological analysis and antibody isolation. Serum antibody titers to SARS-CoV-2 S and N proteins were measured by enzyme-linked immunosorbent assays (ELISA) (**Fig. 1A, B, Table S1**). Serologic analysis demonstrated that serum antibody titers to the N protein were substantially higher than to the S protein in most of the patients. For example, ZD004 and ZD006 had only minimal levels of antibody response to the S protein, while they had much higher antibody titers to the N protein. To be noted, the time from the disease onset to complete recovery from clinical symptoms of COVID19 patient ZD006 was only 9 days (**Table S1**).

**Fig. 1.**
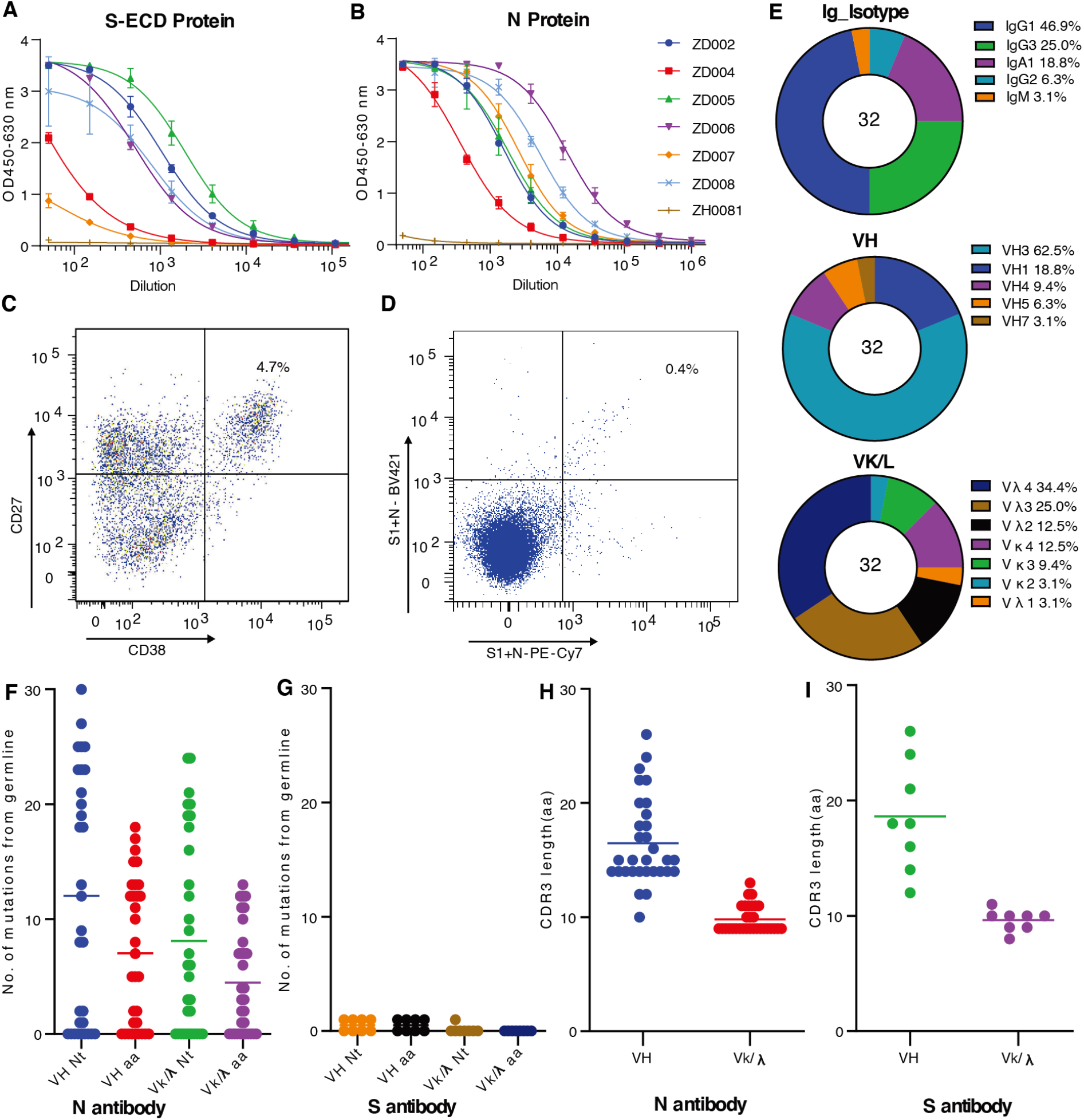
Antibodies acquisition and their characterization. Serum of antibody titers of six SARS-COV-2 convalescent patients to SARS-COV2 S (**A**) and N (**B**) proteins measured by ELISA. Sorting of single plasma cells (**C**) with CD38 and CD27 double positive B cells and single N and S protein-specific memory B cells (**D**) by FACS. (**E**) Percentage of different isotypes, VH and VL gene families of 32 isolated N-reactive antibodies. (**F**) Number of mutations in nucleotides and amino acids in VH and VL (Vκ and V_λ_) of 32 N-reactive antibodies and eight S-reactive antibodies(G). H-CDR3 length of the 32 N-reactive antibodies (**H**) and eight S-reactive antibodies (**I**).

To take advantage of patient ZD006 that was still in the early recovery phase with high possibility of high percentage of antigen-specific plasma cells, single plasma cells (**Fig. 1C**) with phenotype of CD3^-^/CD14^-^/CD16^-^/CD235a^-^/CD19^+^/CD20^low-neg^/CD27^hi^/CD38^hi^, as well as antigen-specific memory B cells with phenotype of CD19^+^/CD27^+^ (**Fig. 1D**) were sorted from PBMC of patient ZD006 by fluorescence activating cell sorter (FACS). To ensure an unbiased assessment, the sorting of antigen-specific memory B cells was carried out with combined probes of both fluorophore-labeled S and N recombinant proteins. Variable region of immunoglobulin (Ig) heavy- and light-chain gene segment (V_H_ and V_L_) pairs from the sorted single cells were amplified by RT-PCR, sequenced, annotated and expressed as recombinant mAbs using the methods as described previously(*28*). Recombinant mAbs were screened against SARS-COV-2 S and N proteins. In total, we identified 32 mAbs reacted with SARS-COV2 N protein including 20 mAbs from plasma cells, and 12 mAbs from memory B cells (**Table S2**). We found that IgG1 is the predominant isotype at 46.9% followed by IgG3 (25.0%), IgA (18.8%), IgG2 (6.3%) and IgM (3.1) (**Fig.1E**). V_H_ gene family usage in SARs-COV2 N protein-reactive antibodies was 18.8% V_H1_, 62.5% V_H3_, 9.4% V_H4_, 6.2% V_H5_ and 3.1% V_H7_, respectively (**Fig. 1F**), which was similar to the distribution of V_H_ families collected in the NCBI database. Nine of 32 SARS-COV-2 N protein-reactive antibodies had no mutation from their germline V_H_ and V_H_ gene segments (**Fig. 1F, Table S2**). Average mutation frequency of the remaining mutated antibodies was 5.3 % (+/-3.6%) in V_H_ and 3.5% (+/-2.7%) in V_L_.

In consistent with the lower serum antibody titers to SARS-COV-2 S protein, we identified only eight SARS-COV-2 S protein-reactive mAbs including 5 antibodies from plasma cells and three antibodies from memory B cells. V_H_ gene segment of the S protein-reactive antibodies had either no mutation (6/8) or minimal mutation (1/300) (**Fig.1G**). There were no significant differences in complementarity-determining region 3 (CDR3) length in amino acid residues between the N- (**Fig.1H**) and S-reactive antibodies (**Fig.1I**).

Approximately a quarter portion of antibodies directed to the N protein (**Fig.1F**) and almost all of antibodies to the S protein that had no mutation or minimal mutations from their germlines (**Fig.1G**) reflected as primary antibody response similar to other typical primary viral infections. However, relatively high V_H_ mutation frequencies (mean of 5.7%) of the majority antibodies to the N proteins were more similar to mutation frequencies of antibodies from the secondary responses to influenza vaccination reported previously. Although patient ZD006 was hospitalized for only nine days after the first appearance of COVID-19 symptoms, the patient has high serum antibody titers and the majority of the isolated N-reactive antibodies have high mutation frequency, whereas the S-directed antibodies have no mutation or minimal mutation. These results reflect much stronger antigen stimulation to the host driven by SARS-COV2 N protein than by the S protein.

### Binding characterizations of anti-N mAbs

To determine the antigenic targets by the N-reactive antibodies, we next analyzed the binding activities by ELISA with variant constructs of the N protein (N-FL: 1-419; N-NTD: 41-174; N-CTD: 250-364) (**Fig. 2A**). Among 32 mAbs binding to NFL; 13 antibodies bound to N-NTD; one antibody bound to N-CTD (**Fig. 2B**). Total of nine antibodies including one antibody (nCoV400) recognizing N-CTD, seven mAbs binding N-NTD (nCoV396, nCoV416, nCoV424, nCoV425, nCoV433, nCoV454, nCoV457) and one mAb (nCoV402) binding only to NFL but not to the other variant N proteins were chosen as representatives for further study. Purified antibodies were confirmed to bind the NFL protein by ELISA (**Fig. 2C**). Affinity of these antibodies to the NFL protein was measured by surface plasmon resonance (SPR) (**Fig. 2D**). In an effort to further characterize the function and structure relationship, three antibodies nCoV396, nCOV416 and nCOV457 were selected for production of recombinant Fab antibodies based on their unique characters. MAb nCoV396 has V_H_ mutation frequency of 2.8%, but high binding affinity with KD of 1.02 nM (**Fig. 2D**) to the N protein. MAbs nCOV416 and nCOV457 have high V_H_ mutation at 11.1% and 8.7%, respectively, and have binding affinity to N protein with KD of 7.26 nM and 12.6 nM (**Fig.2D**, **Table S3**).

**Fig. 2.**
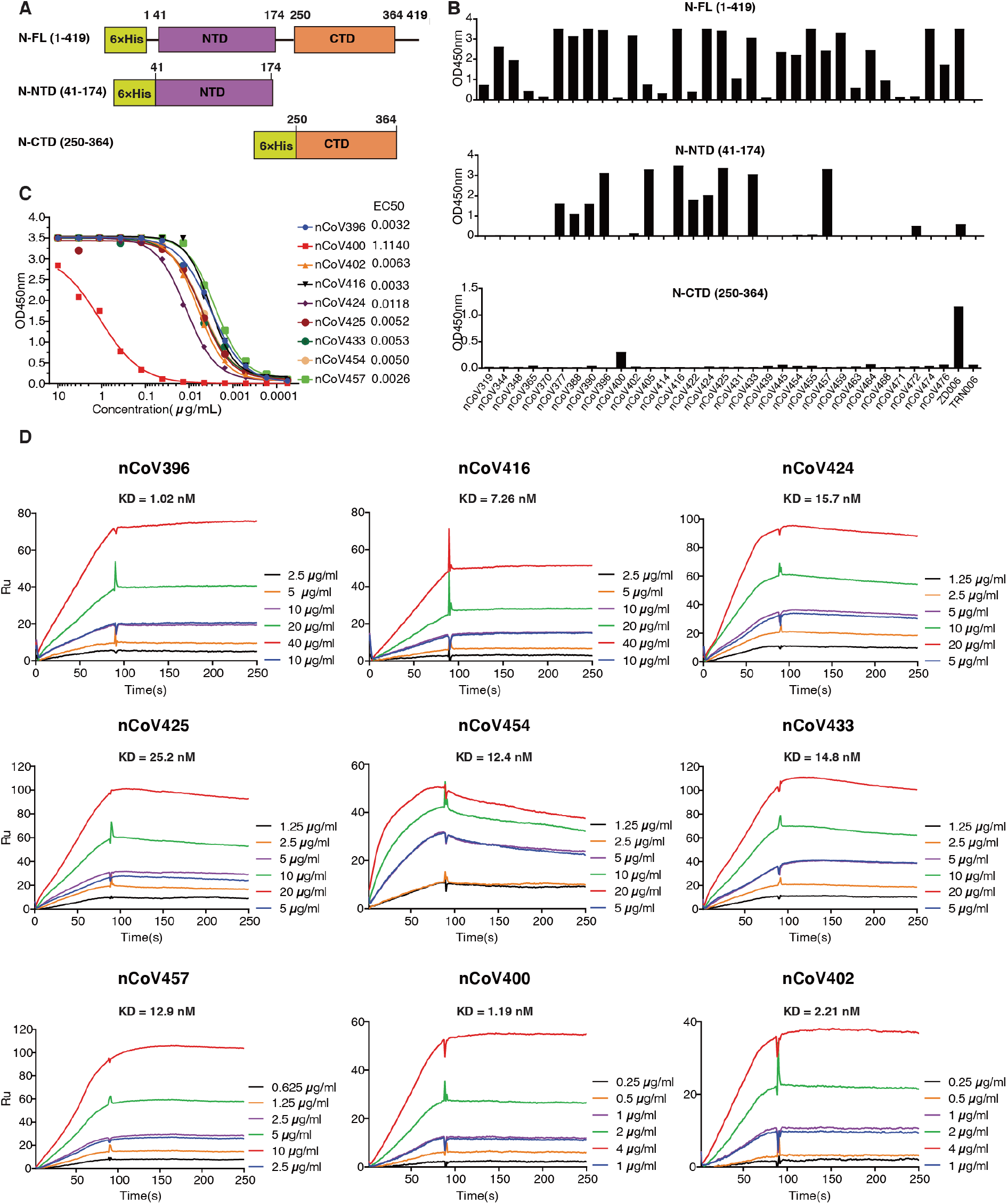
Reactivity and affinity of the isolated antibodies to the N protein antigens. (**A**) Schematic presentation of SARS-COV2 N protein and two variant forms. (**B**) Antibodies expressed in 293 cells transfected were evaluated for binding to the N-FL, N-NTD and N-CTD by ELISA. Plasma from the patient ZD006 and an irrelevant mAb TRN006 were used as positive control and negative control, respectively. (**C**) Ability of nine purified antibodies to the N-FL protein was determined by ELISA. (**D**) Binding affinity of nine selected antibodies to N protein were measured by SPR. KD were shown above the individual plots.

### Complex structure of mAb with N-NTD

To investigate the molecular interaction mechanism of mAb nCoV396 with N protein, we next solved the complex structure of SARS-CoV-2 N protein NTD (N-NTD) with nCoV396 Fab fragments (nCoV396Fab) at 2.1 Å resolution by X-ray crystallography. The final structure is fitted with visible electron density spanning residues 49-173 (SARS-CoV-2 N-NTD), 1-220 (nCoV396Fab, the heavy chain of Fab fragments), and 1-213 (nCoV396Fab, the light chain of Fab fragments, except residues ranged 136-141), respectively. The complete statistics for data collection, phasing, and refinement are presented in **Table S4**.

With the help of the high-resolution structure, we were able to designate all complementarity determining regions (CDRs) in the nCoV396Fab as L-CDR1 (light chain CDR1, residues 23-32), L-CDR2 (light chain CDR2, residues 51-54), L-CDR3 (light chain CDR3, residues 94-100), H-CDR1 (heavy chain CDR1, residues 26-33), H-CDR2 (heavy chain CDR2, residues 51-57), and H-CDR3 (heavy chain CDR3, residues 99-108). Among them, we identified the interaction interface between N-NTD and L-CDR1, L-CDR3, H-CDR1, H-CDR2, H-CDR3 of nCoV396Fab with unambiguous electron density map (**Fig. 3A**, **Fig. S1A**).

**Fig. 3.**
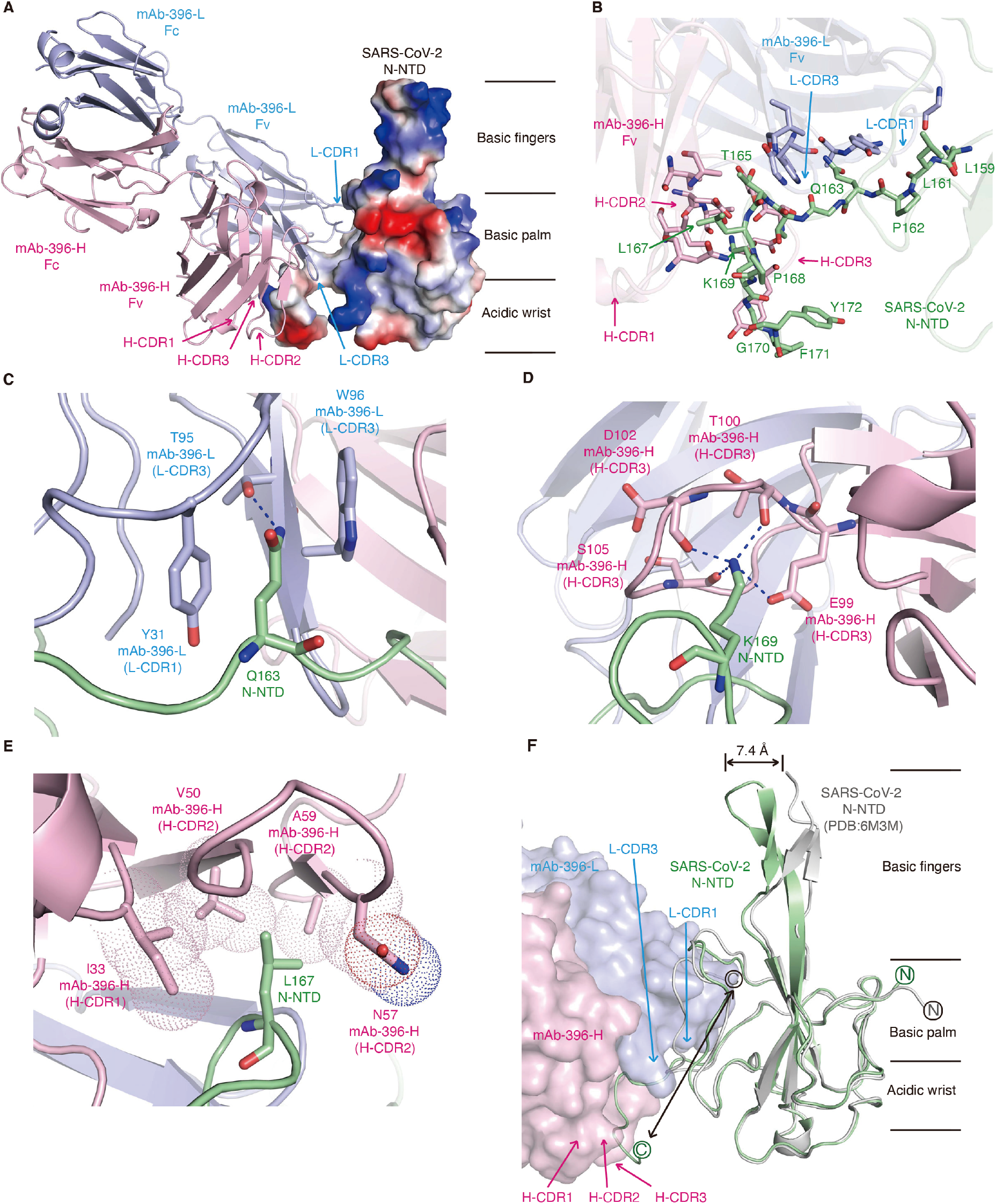
Complex structure of mAb nCoV396 with SARS-CoV-2 N-NTD. (**A**) Overall structure of the mAb nCoV396 - SARS-CoV-2 N-NTD complex. The light chain (pink) and heavy chain (blue) of mAb nCoV396 are illustrated with ribbon representation. SARS-CoV-2 N-NTD is illustrated with electrostatics surface, in which blue denotes positive charge potential while red indicates negative charge potential. (**B**) The N-NTD epitope recognized by mAb nCoV396. The interacting residues of N-NTD and nCoV396 is highlighted with stick representation. Recognition of Q163 (**C**), K169(**D**) and L167 (**E**) in N-NTD by mAb nCoV396. Dash blue line represents hydrogen bond. Hydrophobic interactions are illustrated with dot representation. (**F**) Conformational changes of N-NTD upon the mAb nCoV396 binding. Apo structure of N-NTD is colored with grey. Antibody bound N-NTD is colored with green. N-terminal and C-terminal of the N-NTD is labeled with circle characters. mAb nCoV396 is illustrated with surface representation. All figures were prepared by Pymol.

The interacting CDRs pinch the C-terminal tail of SARS-CoV-2 N-NTD (residues range from 159 to 172), with extensive binding contacts of 1079 Å^2^ burying surface area (**Table S5**). Light chain L-CDR1 and L-CDR3 of nCoV396Fab interact with residues ranging from 159-163 of N-NTD via numerous hydrophilic and hydrophobic contacts (**Fig. 3B, Fig. S1B**). Of note, SARS-CoV-2 N-NTD residue Q163 is recognized by L-CDR3 residue T95 via a hydrogen bond, simultaneously stacking with L-CDR3 residue W96 and L-CDR1 residue Y31 (**Fig. 3C**). Besides, a network of interactions from heavy chain H-CDR2, H-CDR3 of nCoV396Fab to residues 165-172 of N-NTD suggests that SARS-CoV-2 N-NTD conservative residue K169 has a critical role in nCoV396 antibody binding. The K169 is recognized via hydrogen bonds with residues E99 δ-carboxyl group and T100, D102, S105 main-chain carbonyl groups inside the H-CDR3 of nCoV396Fab (**Fig. 3D**). Besides, SARS-CoV-2 N-NTD L167 also interacts with I33, V50, N57, and A59 of H-CDR1 and H-CDR2 of nCoV396Fab through hydrophobic interactions (**Fig. 3E**). Interestingly, all three residues (Q163, L167, and K169) of SARS-CoV-2 N-NTD are relatively conserved in the highly pathogenic betacoronavirus N protein (**Fig. S2B**), which implicated that the nCoV396 may cross-interact with SARS-CoV N protein or MERS-CoV N protein. Indeed, the binding affinities measured by SPR analysis demonstrate that nCoV396 interacts to SARS-CoV N protein and MERS-CoV N protein with KD of 7.4 nM (**Fig. S2B, C**).

To discover the conformational changes between the SARS-CoV-2 N-NTD apo-state with the antibody-bound state, we next superimposed the complex structure with the N-NTD structure (PDB:6M3M)(*17*). The superimposition result suggests that the C-terminal tail of SARS-CoV-2 N-NTD unfold from the basic palm region upon the nCoV396Fab binding (**Fig. 3F**), which likely contributes to allosteric regulation of normal full-length N protein’s function. Additionally, nCoV396Fab binding results in a 7.4 Å movement of the β-finger region outward from the RNA binding pocket, which may enlarge the RNA binding pocket of the N protein (**Fig. 3F**).

To sum up, our crystal structural data demonstrated that the human mAb nCoV396 recognizes the SARS-CoV-2 N protein via a pinching model, resulting in a dramatic conformational change of residues ranged from 159 to 172, which is the linker region of N-NTD connected with other domains.

### MAb curbs N-induced complement activation

Although a recent study suggests that complement cascade is hyperactive by N protein in lungs of COVID-19 patients *via* lectin pathway(*15*), it is unclear how to develop a virus-free and effective system for analyzing the role of SARS-CoV-2 N protein on complement hyper-activation. To this end, we developed a clinical autoimmune disease serum-based protease enzymatic approach to assess complement activation level in the presence of SARS-CoV-2 N protein. Since complement activation initiated by lectin pathway is featured with MASP-2 proteases by specific activity for cleaving complement component 2 and 4 (C2 and C4)(*29*), we designed a complement component 2 (C2) internal quenched fluorescent peptide-based analysis route for *ex vivo* complement hyper-activation (**Fig. 4A**). Briefly, serum was collected from peripheral blood of the volunteers with autoimmune disease, which contains necessary components for complement activations characterized by elevated levels of C3 value (**Table S6**). Next, we collected the fluorescence signal from cleaved C2 synthetic peptide substrates (2Abz-SLGRKIQI-Lys(Dnp)-NH_2_) in reaction mixtures containing autoimmune disease serum, via in the absence or presence of SARS-CoV-2 N protein with or without mAb nCoV396. The initial reaction rate (*v*_0_) was estimated at a single concentration of individual sera from duplicate measurements over a range of substrate concentrations. The steady-state reaction constants *V*_max_ (maximal velocity) and K_m_ (Michaelis constant) were determined for comparisons (**Fig. 4A**).

**Fig. 4.**
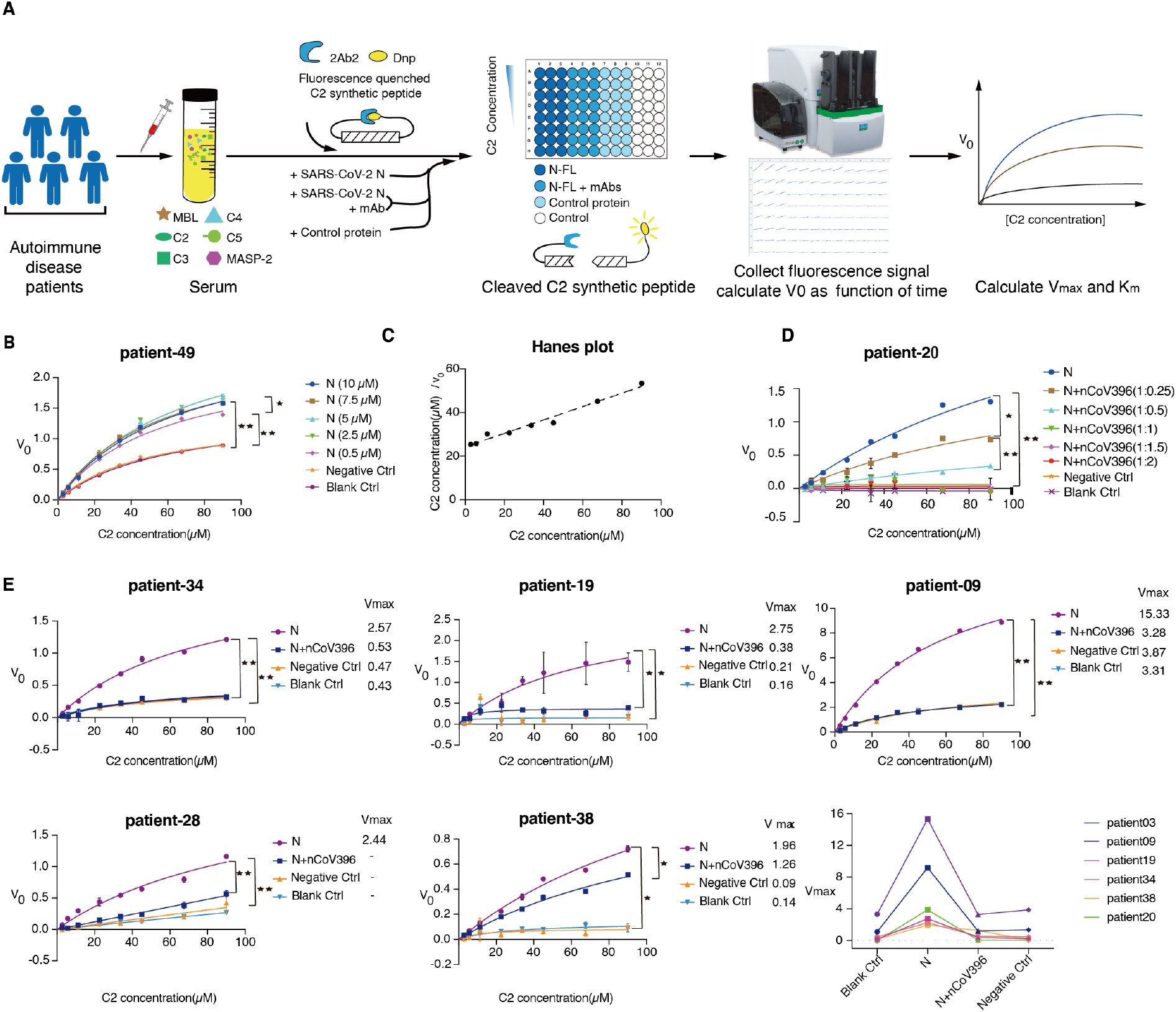
Antibody nCoV396 compromise SARS-CoV-2 N protein induced complement hyper-activation. (**A**) Flow scheme of SARS-CoV-2 N protein and nCoV396 influent the protease activity of MASP-2 in serum of autoimmune disease patients. The Michaelis-Menten curve shows the effect of increasing N protein concentration(**B**) and antibody concentration(**D**) on the substrate C2 cleavage of MAPS2 in serum of patient-49 and patient-20. (**C**) A Hanes plot where C2 concentration/V_0_ is plotted against C2 concentration of adding 5 μM N protein. (**E**) MAb nCoV396 inhibits N protein induced excessive cleavage of C2 in serum of six autoimmune disease patients and last panel shows a summary of Vmax for all patients. Negative control (Negative Ctrl) and blank control (Blank Ctrl) represent reactions containing BSA instead of N or N and mAb, and without exogenous protein, respectively. The mean values and SDs of three technical replicates are shown. P values: *P < 0.05; **P < 0.01; “-” means that the kinetics did not conform to Michaelis-Menten kinetics.

As shown in **Fig. 4B**, the calculated *V*_max_ of reactions without any other exogenous proteins is 1.49 RU·s^-1^. Additions of SARS-CoV-2 N protein (concentrations ranged 0.5 μM to 10 μM) in the reactions remarkably elevate the *V*_max_ up to 2 folds, ranged from 2.37 ~ 3.02 RU·s^-1^. Similarly, additions of SARS-CoV-2 N protein lead to approximate 1.8 folds increasing of the *V*_max_/K_m_ values, which suggested that the specificity constant (K_cat_/K_m_) of MASP-2 to substrates is increased in the presence of viral N protein as the enzyme concentrations are equivalent among the reactions (**Table S7 - S8**). To confirm the kinetic analyses, Hanes plots ([S]/V versus [S]) were also drawn and found to be linear (**Fig. 4C**). Therefore, the additions of SARS-CoV-2 N protein do not change the single substrate binding site characterization of the enzymatic reactions. To assess the suppression ability of nCoV396 to the SARS-CoV-2 N protein-induced complement hyper-activation function, we next conducted the complement hyper-activation analysis in serial N protein: nCoV396 ratios. As shown in **Fig. 4D**, the addition of N protein elevates *V*_max_ value up to 40-folds (1:0 ratio), whereas the additions of antibody nCoV396 decline the *V*_max_ in a dose-depended manner (**Table S9**). To further validate the function of nCoV396, we next perform complement hyper-activation analysis in other five serum samples from autoimmune disease donors. Consistently, the *V*_max_ of reactions are boosted in the presence of N protein in all samples, while declined in the presence of mAb nCoV396 together with N protein (**Fig. 4E)**. In conclusion, these results demonstrate that SARS-CoV-2 N protein is capable of inducing the complement hyper-activations *ex vivo*, not only by facilitating the maximal velocity of MASP-2 catalytic activity, but also enhancing the substrate binding specificity in the reactions. The N-directed mAb nCoV396 specifically compromises the SARS-CoV-2 N protein-induced complement hyper-activation within clinical serum samples.

## Discussion

From a quickly recovered COVID-19 patient, we isolated 32 mAbs specifically targeting to SARS-CoV-2 N protein. The binding affinity of mAbs ranged from 1 nM to 25 nM, comparable with mature spike protein-directed antibodies(*20, 23–27*) and the other mature antibodies identified during acute infections(*30, 31*). Characteristics of the isolated N-reactive mAbs are different from the isolated S-reactive mAbs in the early recovery COVID-19 patients suggested that sampling time is pivotal for identifying differential immune responses to different SARS-CoV-2 viral proteins.

The crystal structure of nCoV396 bound to SARS-CoV-2 N-NTD elucidates the interaction mechanism of the complex between the first reported N protein-directed human mAb and its targeted N protein. Three conservative amino acids (Q163, L167, K169) in N protein are responsible for nCoV396 recognition, which provided a clue of cross-reactivity to SARS-CoV or MERS-CoV N protein for nCoV396. Intriguingly, the nCoV396 binding of SARS-CoV-2 N-NTD undergoes several conformational changes, resulting in a change in N-NTD RNA binding pocket enlargement and partial unfolding of basic palm region. More importantly, this conformational change occurs in the C-terminal tail of the N-NTD, which may alter the positioning of individual domains in context of full-length protein and lead to a potential allosteric effect for protein functions.

Complement is one of the first lines of defense in innate immunity and is essential for cellular integrity, tissue homeostasis, and modifying the adaptive immune response(*32*). Emerging evidence suggests that the complement system plays a vital role in a subset of COVID-19 critical patients, with features of atypical acute respiratory distress syndrome, disseminated intravascular coagulation, and multiple organs failure(*9, 10, 33*). A few pieces of evidence show that highly pathogenic coronavirus (i.e., SARS-CoV-2 and SARS-CoV) N protein is involved in the initiated MASP-2 dependent complement activation(*15, 34*). Encouragingly, COVID-19 critical patients treated with complement inhibitors, including small molecules to complement component C3 (AMY-101) and antibody targeting to complement component C5 (Eculizumab), show remarkable therapeutic outcomes(*15*). Currently, there are 11 clinical trials relative to targeting the complement pathway (https://clinicaltrials.gov). In order to avoid adverse effects of human complement component targeting therapy, a viral protein-specific approach is warranted. The antibody nCoV396 isolated from COVID-19 convalescents is an excellent potential candidate with high binding affinity to N protein and high potency to inhibit the complement hyper-activation. As revealed by atomic structural information, the binding may allosterically change the full-length N protein conformation. To determine the role of nCoV396 in the suppression of complement hyper-activation, we monitor the MASP-2 protease activity based on its specific fluorescent quenched C2 substrate in serums from autoimmune disease patients. The complete complement components in sera of patients with autoimmune disorders allow us to monitor the activating effects of SARS-CoV-2 N protein and its specific mAbs. Although we cannot calculate the other steady-state enzymatic reaction constants as the precisely concentration of MASP-2 in serum is unknown, we identified the *V*_max_ of the specific C2 substrate for the enzymatic reaction. We demonstrated that SARS-CoV-2 N protein elevated the *V*_max_ of the reaction, up to 40 folds, in serum of all 7 individuals tested, while nCoV396 effectively suppress *V*_max_ of the reaction mixture. These results indicated that the autoimmune disease patient serum-based complement activation analysis is a virus-free and effective method for examining complement activation mediated by coronavirus N protein.

Although precise interaction of SARS-CoV-2 N protein with MASP-2 remains to be elucidated, our work defined the region on the SARS-CoV-2 N protein recognized by mAb nCoV396 that plays an important role on complement hyper-activation, and indicates that human mAbs from the convalescents could be a promising potential therapeutic candidate for the treatment of COVID-19.

## Acknowledgments

We thank the staffs of the BL18U/19U/17U beamlines at SSRF for their help with the X-ray diffraction data screening and collections. We thank Junlang Liang, Tong Liu, Nan Li, Xiaoli Wang, Zhenxing Jia, and Jiaqi Li from Zhuhai Trinomab Biotechnology Co., Ltd. for technical assistants of mAbs isolation, production and characterization.

## Author contributions

S. C., H. S., F. X. and H-X. L. contributed the conception of the study and established the construction of the article. S. C. and H-X. L. designed the experiments and wrote the manuscript. S. K., M. Y., S. H. contributed to protein purification and crystallization, in vitro protein-protein interaction analysis, and complement activation analysis. Y. W. contributed to mAbs isolation, in vitro protein-protein interaction analysis. S. C., S. K. M. Y., and S. H. performed structural determination and validation. S. C., S. K., Y. W. drew figures. X. C., Y. C., Q. C., Z. Z., Z. Z., Z. H., X. H., H. S., W. Z., and H. H. contributed to interpretation of data. Z. H., J. L., G. J., and F. X. contributed to clinical samples collections. S.K., M.Y., S. H., Y.W. contributed equally to this work.

## Data and materials availability

The structure in this paper is deposited to the Protein Data Bank with 7CR5 access code.

